# Peptide therapeutic leads for multi-target inhibition of inflammatory cytokines in Inflammatory Bowel Disease - computational design and in-vitro validation

**DOI:** 10.1101/2024.08.27.609829

**Authors:** Tomer Tsaban, Gali Kariv-Attias, Alisa Khramushin, Ofer Gover, Zvi Hayouka, Ora Schueler-Furman, Betty Schwartz

**Author notes:** Equal contributions (Tomer Tsaban, Gali Kariv-Attias).

## Abstract

Inflammatory Bowel Disease (IBD) are chronic and recurrent inflammatory disorders affecting the gastrointestinal tract, characterized by the involvement of numerous pro-inflammatory cytokines. These conditions profoundly impact both immune system dynamics and intestinal tissue integrity. Current therapeutic approaches predominantly rely on monoclonal antibodies, and frequently encounter limitations such as non-responsiveness, loss of efficacy over time, immunogenicity, adverse effects, and substantial cost. Consequently, there is a critical need for novel, targeted anti-inflammatory strategies. We present the computational structure guided design of peptidic inhibitors aimed at attenuating the activity of pivotal pro-inflammatory cytokines implicated in IBD pathogenesis, namely TNFα, IL-1β, and IL-6. These peptides were engineered to disrupt specific cytokine - receptor interactions, to block the release of pro-inflammatory cytokines. We structurally characterized key features in the studied interactions and used these to guide two computational design strategies, one based on the identification of dominant segments using our PeptiDerive approach, and one based on complementing fragments detected using our PatchMAN protocol. The designed peptides were synthesized and their efficacy was validated on Caco-2 intestinal epithelial cells and THP-1 macrophages, representative of the epithelial and immunological alterations typical of active IBD. The majority of the novel peptides effectively suppressed release of pro-inflammatory cytokines by both macrophages and intestinal epithelial cells, thereby reducing the risk of inflammation. This study underscores the efficacy of a rational design approach rooted in structural insights into inflammatory signaling complexes. Our findings demonstrate the potential of targeting key cytokines and receptor interaction with designed peptides as a promising therapeutic avenue for managing IBD and other inflammatory disorders.

## Introduction

The term Inflammatory Bowel Disease (IBD) describes chronic inflammatory conditions primarily affecting the gastrointestinal tract, specifically the colon, small intestine, or both. One principal form of IBD is Ulcerative Colitis (UC), characterized by inflammation localized to the colon. The second, Crohn’s disease (CD), can affect any segment of the gastrointestinal tract in a non-continuous manner (mouth to the anus). In recent years, there has been a notable global increase in the prevalence of IBD, particularly in newly industrialized nations. Estimates suggest that by 2017 the number of individuals affected worldwide surpassed 7 million [^1^]. This rise underscores the growing public health challenge posed by IBD, necessitating enhanced efforts in research, diagnosis, and management strategies to mitigate its impact on affected populations.

The etiology and mechanisms leading to the formation and progression of IBD are still only partially understood. Genetics, environmental exposure to food and pollutants, and stress factors, seem to play a major role. Multiple studies in the past 20 years have highlighted the complex cytokine network involved in initiating the disease and mediating its progression. Prominent among these molecular factors are Tumor necrosis factor-alpha (TNF-α), IL6, IL1-β, and IL-12, all playing a critical role in both UC and CD [^2^ ^3^].

TNF-α is a key mediator in the inflammatory processes of the intestine and plays a crucial role in the pathogenesis of IBD. Elevated serum levels of TNF-α are frequently observed in patients with IBD. Furthermore, TNF-α is instrumental in driving inflammation through the induction of other pro-inflammatory cytokines, such as interleukin-1 beta (IL-1β) and interleukin-6 (IL-6) [^4^]. During active colonic Inflammation, TNF-α triggers other immune cells, promotes chemotaxis and enhances intracellular pathogen destruction by releasing toxic oxygen radicals and nitric oxide [^5^].

IL-6 plays a pivotal role in regulating vascular homeostasis and cellular inflammation, making it a crucial marker of cytokine storms. Evidence demonstrated that numerous pro-inflammatory cytokines and chemokines, including IL-6 are involved in the pathogenesis and development of IBD. Since IL-6 levels are positively associated with the severity and clinical disease activity of IBD patients it is tenable to hypothesize that therapeutic strategies targeting the inhibition of IL-6 during stages of hyperinflammation characterized by dysregulated IL-6 production hold promise for treating IBD [^6^].

IL-1β is a crucial mediator of inflammation and tissue damage in IBD. The equilibrium between IL-1β and its endogenous antagonist, IL-1 receptor antagonist (IL-1Ra), is crucial for the initiation and regulation of inflammatory processes. IL-1β significantly contributes to the induction of pro-inflammatory responses by recruiting and activating immune cells to the intestinal mucosa. Concurrently, IL-1β is involved in disrupting the intestinal barrier and modulating T helper (Th) cell differentiation, particularly influencing Th17 cell development, which is implicated in IBD pathogenesis [^7^]. Furthermore, gut dysbiosis can stimulate immune cells to secrete IL-1β, thereby exacerbating inflammation[^8^ ^9^]. In summary, due to IL-1β’s critical role in driving inflammation and modulating immune responses in IBD, targeting this cytokine or its receptors holds considerable promise as a therapeutic strategy.

Initial mapping of the molecular factors allowed for the development of treatments targeting fundamental IBD cytokines. Anti-TNFα antibodies, introduced in the early 2000’s, gradually replaced the “conventional strategy”, based on non-specific anti-inflammatory treatments (such as corticosteroids). In the following ∼20 years, several more biological treatments were introduced, the vast majority targeting TNFα (Infliximan, Adalimumab, Golimumab) and T-cell integrins (Vedolizumab e.g.), and few targeting IL-12/IL-23. Additionally, clinical trials are ongoing for therapeutics targeting IL-6, IL-18, IL-1 and several others [^10^ ^11^].

While biological treatments have revolutionized the field, a large portion of IBD patients still remains unresponsive to therapy. Among them, a high percentage develops resistance or cannot tolerate adverse effects, varying depending on the specific treatment and patient population. Generally, estimates suggest that approximately 30-40% of IBD patients may not respond to initial biologic therapies or may develop loss of response over time, and adverse effects that lead to discontinuation or intolerance can affect a smaller but significant subset of patients [^12^]. Additionally, a growing body of evidence supports early administration of biologics due to their efficiency profile, and their potential to divert the natural history of the disease [^13^]. Alongside the rise in prevalence, this worsens the ever-growing economic burden of IBD treatment [^13^ ^10^].

Many of these caveats may be attributed to the heavy reliance on monoclonal antibody therapeutics for IBD [^14^ ^10^]. Antibodies offer a straightforward approach for potent inhibition of cytokines and proteins in general. Due to their large size and globular structure, they are biochemically stable and consist of a large potential binding interface and high affinity. However, this makes it hard to administer them pharmacologically. They are also likely to generate immunogenicity which may lead to loss of response or adverse effects. Since IBD has a spectrum of clinical and molecular manifestations, and given antibodies are optimized to interfere with one specific interaction out of a vast, complex inflammation network, some patients are likely to be non-responsive. Additionally, antibodies are harder to express, synthesize, and preserve, making such biologic drugs highly expensive. Hence, low-cost biosimilar agents have been suggested as a promising direction for IBD treatment [^15^].

In recent years, mini proteins, peptides (short protein segments) and peptide mimetics are emerging as promising therapeutic candidates. They combine the specificity of large protein interfaces with the benefits of small sized molecules, including easier production, facilitated cellular transport and lower immunogenicity [^16^ ^17^ ^18^ ^19^ ^20^]. Modulatory peptides are often screened using high-throughput methods (such as phage display), or designed, either computationally or using experimental approaches. The process of computational peptide design may be categorized into three approaches: a) Structure guided design, involving detection and selection of natural structural interaction motifs followed by their optimization; b) De novo design, building totally new potential binders and ranking them; and c) Machine learning (ML) based design using dedicated ML algorithms trained on protein and peptide structural data [^20^].

Such design approaches were recently employed by Huang *et al.* to develop potent pico-molar mini protein modulators that target IL-6 and IL-1 receptors [^21^]. Their strategy involved massive computational sampling of de-novo mini protein scaffolds, docking scaffolds to cytokines yielding millions of structural designs, followed by experimental validation of ∼100k sequences aimed to fold into the mini protein designs. On the experimental front, Hommel *et al.* applied a high-throughput screen of thousands of small molecules to bind IL-1β, identifying a fragment that binds with low μM affinity to a novel binding site [^22^]. For peptide modulators, studies have demonstrated their anti-inflammatory potential [^23^ ^24^], and specifically in IBD [^25^ ^26^]. Jiang *et al.* [^27^ ^28^] manually selected natural motifs from structures of several cytokine interactions (TNFα, IL6, and IL1β) and concatenated the selected peptide candidates into one long chimeric peptide, aimed to potently inhibit three cytokine interactions simultaneously. They demonstrated that it reduces cellular inflammation in gut epithelial cells.

Using several peptides to target multiple core cytokines of the IBD inflammation network is a promising avenue to overcome the aforementioned challenges of non-response and loss of response, and simplifies subsequent trials that test adverse effects such as the immunogenicity of novel compounds. The plethora of computational, energy-based, low resource methods available for the determination of “hot” interaction interfaces can help define interfaces of interest, which in turn can be used for improved natural motif selection for structure guided design, or to filter for relevant potential de novo binders.

In this work we sought to efficiently design a set of peptides aimed at three core cytokines: TNFα, IL1β and IL6, and their interactions with their receptors, which are at the basis of IBD. We utilized computational tools to detect potential binding pockets, stratified interaction residues by their contribution to binding energy and overlaid them with known experimental hotspots. Then, we selected natural interaction motifs using the PeptiDerive protocol previously developed by us [^29^ ^30^] which for each interaction partner derives segments that contribute the most to binding energy, based on a structure of the complex. In cases where no promising natural motifs were identified, we used a more general approach based on the PatchMAN protocol previously developed by us [^31^]. PatchMAN maps short backbone “seeds” extracted from proteins with local structural similarity to interfaces of interest. The identified fragments can be used as starting points for the design of stronger binding peptides. Not only does this remove the dependence on solved structures of interactions which is in some cases a great challenge, it also allows to generate peptides that cover several binding hotspots, even if in the original interaction these were not contacted by nearby sequence positions. Similar approaches have also been suggested by others [^32^ ^33^ ^34^].

To evaluate the effect of our proposed designs on the immunological response typically expressed during IBD, we synthesized the designed peptide and tested them on experimental cell models associated with intestinal inflammation - Caco-2 human intestinal epithelial cells and activated THP-1 macrophage human cells. Our assumption is that the peptides will prevent the cytokine-receptor interaction and reduce the intensity of inflammation by blocking the expression of proinflammatory cytokines and the resulting intestinal tissue damage, thereby inhibiting disease progression. The different peptides were designed to specifically interfere with synergistic effects exerted by TNFα/IL-1β and IL-6 on both intestinal cells and intestinal immune cells, an effect that ultimately amplifies the anti-inflammatory activity of these peptides. Once validated for their individual activity, we will in a future study optimize a combination of these peptides to achieve a multi-pronged drug.

## Results

### Design of peptides targeted to inhibit specific cytokine-receptor interactions: Overview

As a starting point the interacting partners and their relevant interfaces were defined. Using the structure of the complex of the target cytokine and its receptor alongside with experimental evidence collected from the literature, each interaction was mapped for key structural features, including computational alanine scanning and solvent mapping (see Methods). Areas covering these were defined as its core structural components.

We then employed two peptide design strategies (**Figure 1**): **PeptiDerive Strategy A** (**Figure 1B**) included mapping the segment that contributes most of the energy in binding to the partner, and deriving peptides from these segments (using our PeptiDerive approach [^30^ ^29^], see Methods). These peptide segments (referred to as “natural structural motifs” in [^20^], as described in the introduction) were scored according to their computed energy and their overlap (recapitulation) of the key features that characterize the interaction. Additionally, each candidate was manually examined and evaluated for its potential to maintain structural integrity out of the derivation context.

**Figure 1:**
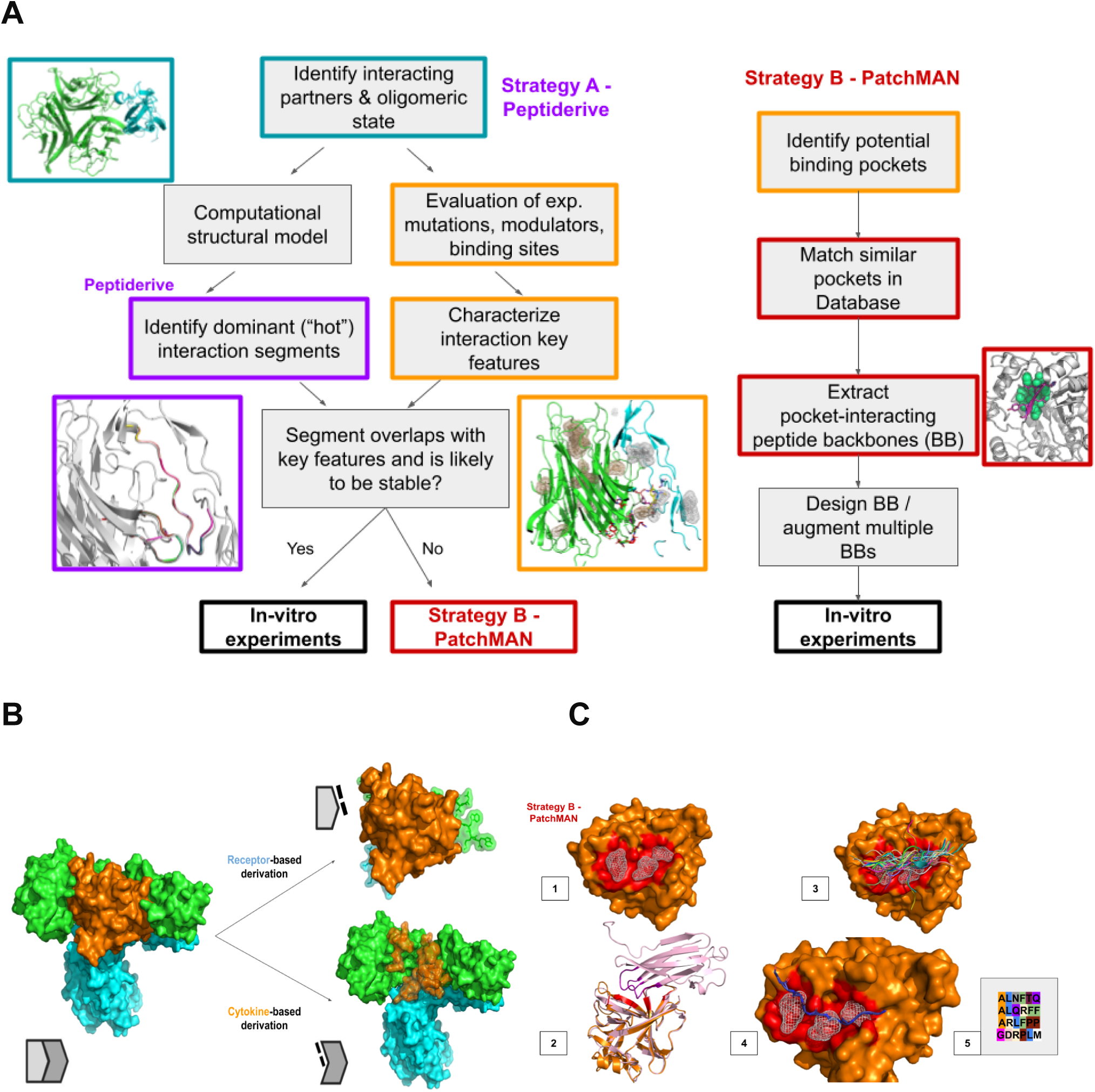
(A) Overview of strategies used to design inhibitory peptides. PeptiDerive Strategy A (left, **B** below) involves the determination of dominant peptide segments (natural structural motifs) in a given structure of an interaction (see **B**), and selecting those that contribute most to binding energy and target known binding pockets. If no such peptides are found, Strategy B is used. PatchMAN Strategy B (right, **C** below) involves the mapping of the receptor surface with complementing peptide backbones, which are extracted from similar structures (see **C**). These backbones are then redesigned to bind optimally to the receptor surface (see Text for more details). **(B) Design strategy A: Derive dominant segments using PeptiDerive** (illustration). Left: a cytokine-receptor complex, (orange and green-cyan respectively), showing the surfaces of the partners. Peptides are derived from interacting segments, from each of the partners (Top right: green and cyan peptides derived from the receptor units, interacting with the orange cytokine; Bottom right: orange peptides derived from the cytokine, interacting with the receptor units). **(C) Design strategy B: Determine peptide fragments using PatchMAN** (illustration). **1. Define binding pockets** (using computational solvent mapping with the FTmap protocol - white mesh) on target (orange surface, binding pocket in red). **2. Detect peptide backbone fragments that can interact with the pocket surface** (using PatchMAN): The patch covering pocket-forming residues (in red) is searched using MASTER to identify matching structures (example match in pink), from which short peptide backbones that interact with the area matching the query were extracted to serve as scaffolds (in purple). **3. Ensemble of peptide scaffolds** collected from all structural matches **4. Select scaffold(s) that cover the pocket(s) of interest** (wheat and pale blue) **5. Design tightly binding peptide sequence** (using Rosetta design).

For interactions with no lead peptide candidates, we proceeded to **PatchMAN Strategy B (Figure 1C)**, in which peptide backbone conformations that complement a given surface patch are extracted from similar structures (using our PatchMAN approach [^31^], see Methods), and their sequence is redesigned to optimally fit the receptor (using Rosetta FlexPepDesign, see Methods).

### Exploring the lead peptides activity

The designed peptides were synthesized and experimentally validated in three different in vitro set-ups:

**(1) Impact of the designed peptides on the secretion of IL-8 in Caco-2 epithelial cells.**IL-8 is a potent chemokine crucial for recruiting neutrophils and granulocytes to sites of tissue damage [^35^] that is produced by various cell types including blood monocytes, fibroblasts, endothelial cells, and epithelial cells. There is established evidence indicating elevated IL-8 production in the tissue of patients with IBD, particularly in those with Ulcerative Colitis (UC), compared to healthy individuals [^36^]. Consequently, there is active development of therapeutics aimed at targeting IL-8 as a potential treatment strategy for these conditions. Our experiments assessed the effect of each peptide on IL-8 release from Caco-2 cells that were either primed with recombinant TNFα (rTNFα) or unstimulated (as a negative control). **(2) Change in TNFα secretion in THP-1 macrophage cells after treatment with the designed peptides:** The human monocytic cell line THP-1 is an experimental macrophage cell model associated with intestinal inflammation [^37^]. THP-1 cells were initially treated to induce differentiation (M0) by treating them with PMA. M0-THP-1 cells were then classically activated to type 1 (M1) in vitro using the bacterial cell wall component LPS. We tested the effect of each peptide on the release of the key markers TNFα, and IL-1β. **(3) For selected lead peptides that showed promising effects in the secretion experiments we also measured the change in mRNA expression levels of selected cytokines** to further validate that the inhibition of the secretion was due to the blockage of the signal transduction that leads to their expression in CaCo2 and THP-1 cells. To evaluate the putative cytotoxicity of the different peptides, we first performed an MTT assay and verified that the designed peptides have no cytotoxic effect up to the highest concentration that was tested (see **Methods, Supplementary Figures S1 and S2**). To achieve maximal stimulation, we evaluated different rTNFα concentrations and incubation times (**Supplementary Figure S3A-C**), and chose to use in our further experiments on Caco-2 cells 100 ng/ml TNFα for 12 hours incubation for protein secretion assays and 4 hours incubation for gene expression. We also calibrated the concentration for the TNFα monoclonal antibody as a negative control (mAb) for the chosen concentration for rTNFα (**Supplementary Figure S3D**).

In the following we detail the design process for each of the four cytokines targeted in this study. A summary of the peptides that were synthesized and tested is provided in **Table I** (see Supplementary Table S1 for additional information).

**Table I.**
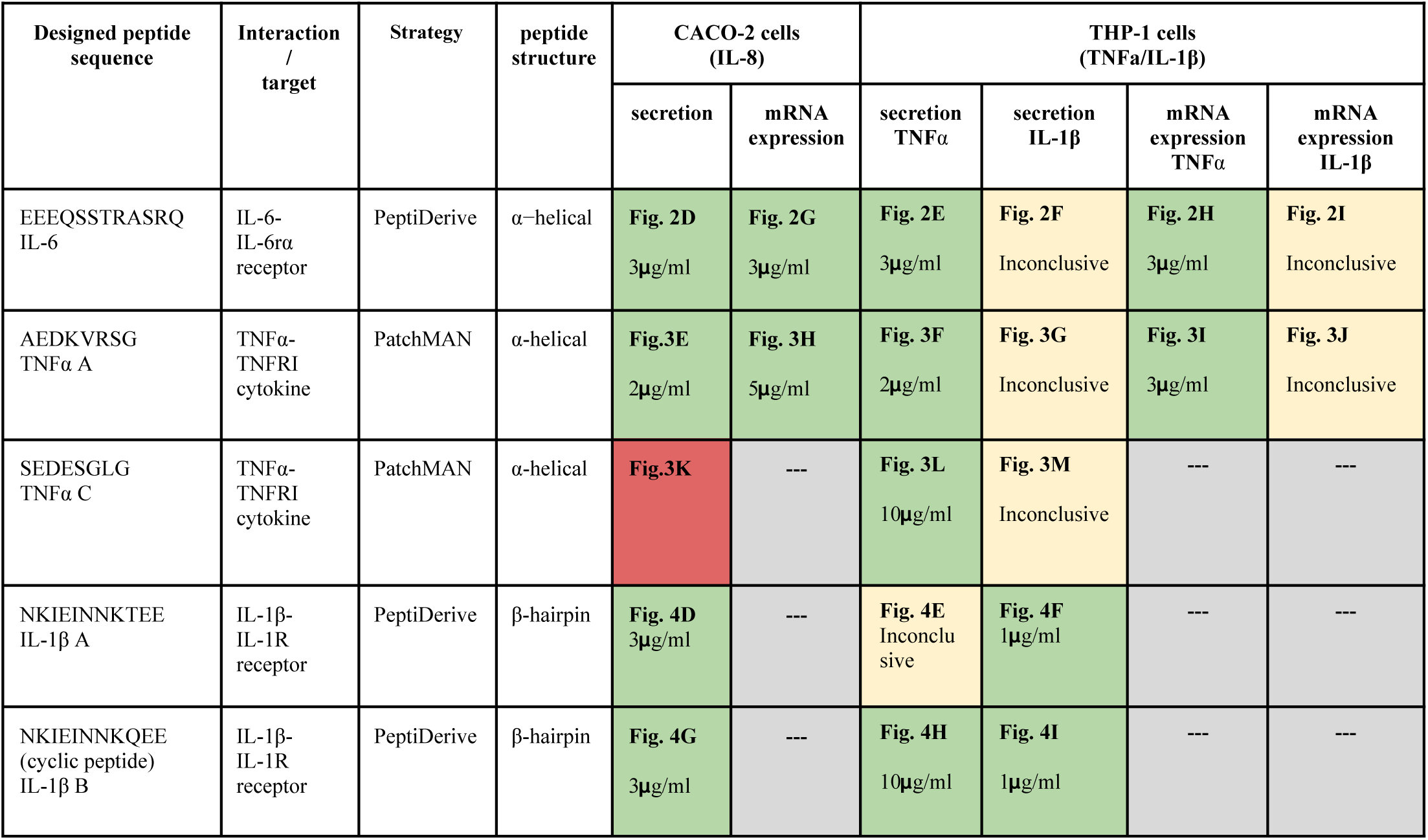
Summary of the designed peptides that were synthesized and characterized in the current study. Cells in green/red/gray/yellow indicate effect/no effect/not measured/inconclusive, respectively, the minimum concentration with significant effect, and the figure with the corresponding results. TNFα B (EAEDARDS) could not be tested due to aggregation.

### Design of α-helical IL-6 inhibitor using PeptiDerive

IL-6 first binds to its receptor IL-6Rα, which positions it to interact with two opposite copies of gp130, leading to the downstream activation of signaling **(Figure 2A)**. We focused on the IL-6 - IL-6Rα interface, where also most key features of the interaction are located: hotspots that were determined experimentally and computationally (using the Rosetta alanine scanning protocol [^38^]), as well as potential binding pockets (determined using computational solvent mapping with FTmap [^39^], see Supplementary Materials for details). We selected an α-helical peptide derived from IL-6 that recovers three experimental hotspots, and interacts with several more **(Figure 2B,C and Supplementary Table S1A for additional details)**. This IL-6 interface designed peptide was synthesized **(EEEQSSTRASRQ)** and we explored its effect in human Caco-2 epithelial cells and in THP-1 human macrophage cells as observed in **(Figure 2D-I)**.

**Figure 2:**
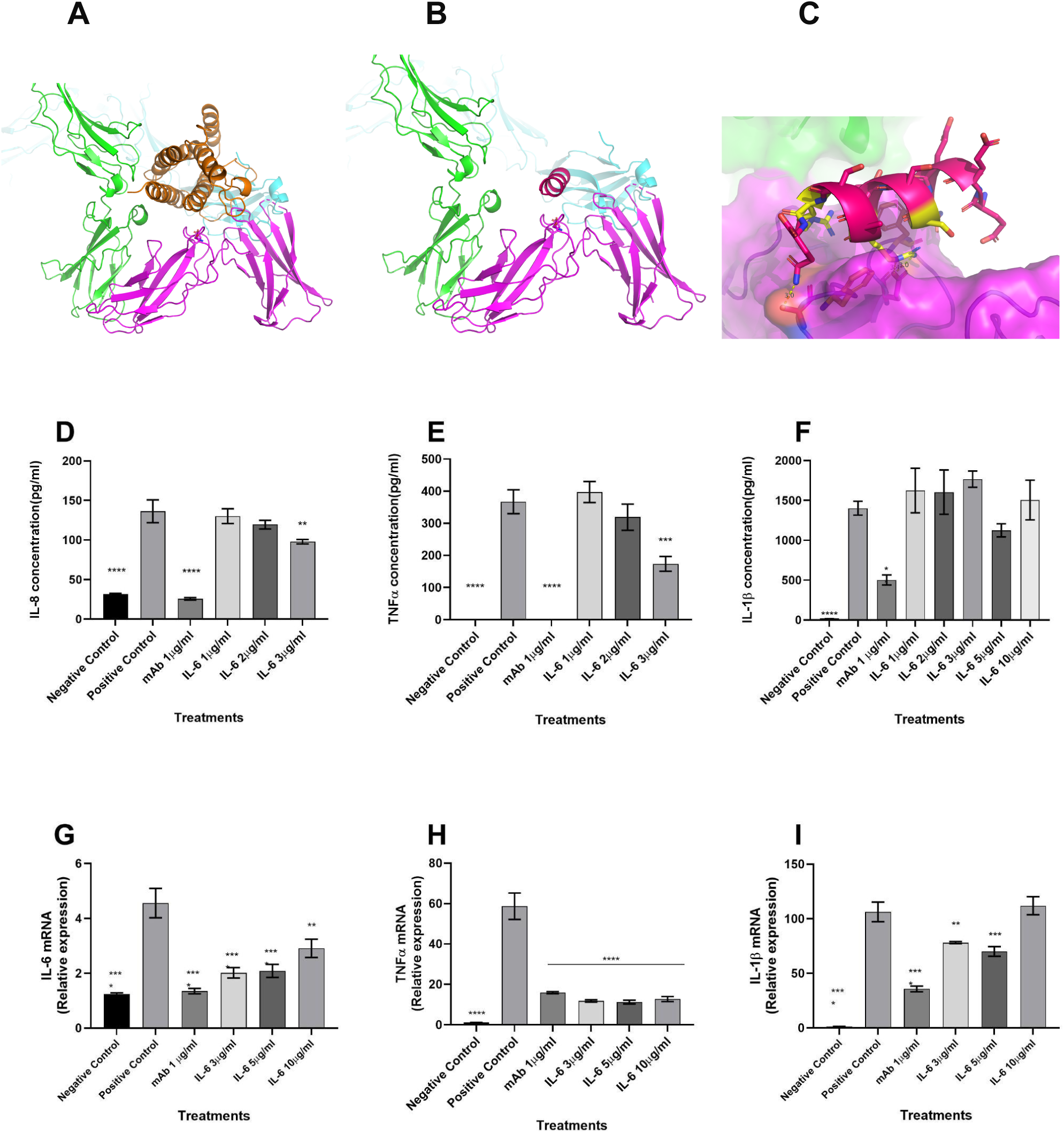
Targeting the interface between IL-6 and IL-6Rα using α-helical peptide. A-C computational design. D-I: Experimental evaluation of effect of addition of increasing concentrations of the designed IL-6 derived peptide (EEEQSSTRASRQ) after 24h on primed Caco-2 intestinal cells and M1-activated THP-1 cells. **(A)** Overview of IL-6 (orange) binding to receptor IL-6Rα (magenta) and two gp130 subunits (green and cyan) (PDB id 1p9m). **(B)** Overview of the designed helical peptide derived from IL-6 binding to the receptor. **(C)** Details of the interaction of the designed peptide with the receptor, hotspot residues colored yellow. **(D)** Caco-2 intestinal epithelial cells were primed with recombinant TNFα (100 ng/ml), and IL-8 release was measured. **(E,F)** THP-1 cells were primed with LPS (10 μg/ml), and release of TNFα **(E)** or IL-1β **(F)** was measured. **(G-I)** Changes in mRNA expression levels of the corresponding cytokines in the corresponding cells. Negative controls represent THP-1 or Caco-2 cells without treatment, and the gold standard inhibition of immune response is included for comparison (1 μg/ml mAb antibody). N=4-6 for all. * pval<0.05, ** pval < 0.005, *** pval < 0.0005, **** pval <0.0001.

### Experimental validation of IL-6 peptide shows its efficiency in reducing inflammatory response

Addition of the IL-6 peptide showed a dosage dependent increasing effect, and at concentrations of 3 μg/ml significantly reduced IL-8 secretion from primed CaCo-2 cells, as well as TNFα secretion from M1-activated THP-1 cells (**Figure 2D,E**). Importantly, the observed effects depend on the addition of TNFα, since the peptide by itself does not affect IL-8 secretion in Caco-2 cells (Data not shown). Interestingly, the inhibitory effect is also reflected in predominant significant downregulation of mRNA expression upon addition of IL-6 derived peptide (**Figure 2G,H**), however, not in a dosage dependent manner. This peptide was ineffective in reducing IL-1β at all concentrations tested (**Figure 2F,I**). Overall, we conclude that the IL-6 derived peptide significantly affects the inflammatory response at the tested concentrations.

### Design of α-helical TNFα inhibitor using PatchMAN

The TNFα cytokine forms a homotrimer that binds 3 monomer receptor units of TNFR1. We focused on one interface formed by two TNFα molecules bound to one TNFR1 molecule **(Figure 3A)**. The key determinants of this interaction involve the membrane distal extracellular part where the TNFα dimer groove contacts the N-terminal CRD1 of the receptor. This region includes mutations reported to impair binding [^40^ ^41,42^], as well hotspots, and several potential binding pockets. This is also reflected by the best scored PeptiDerive results, all focused in the discussed region **(Figures 3A,B**, see Supplementary Materials for details). Hence, this area was our main target.

**Figure 3:**
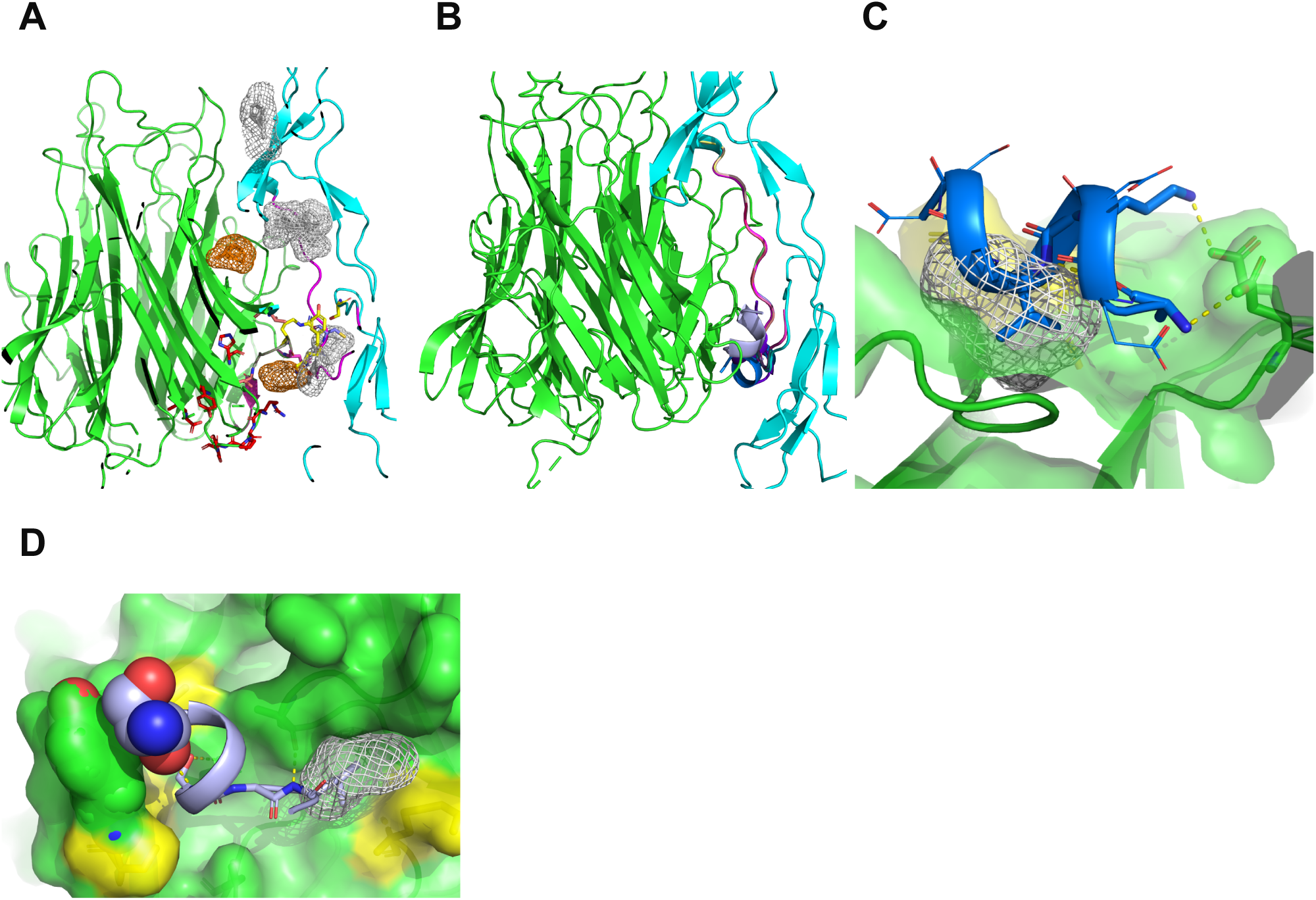
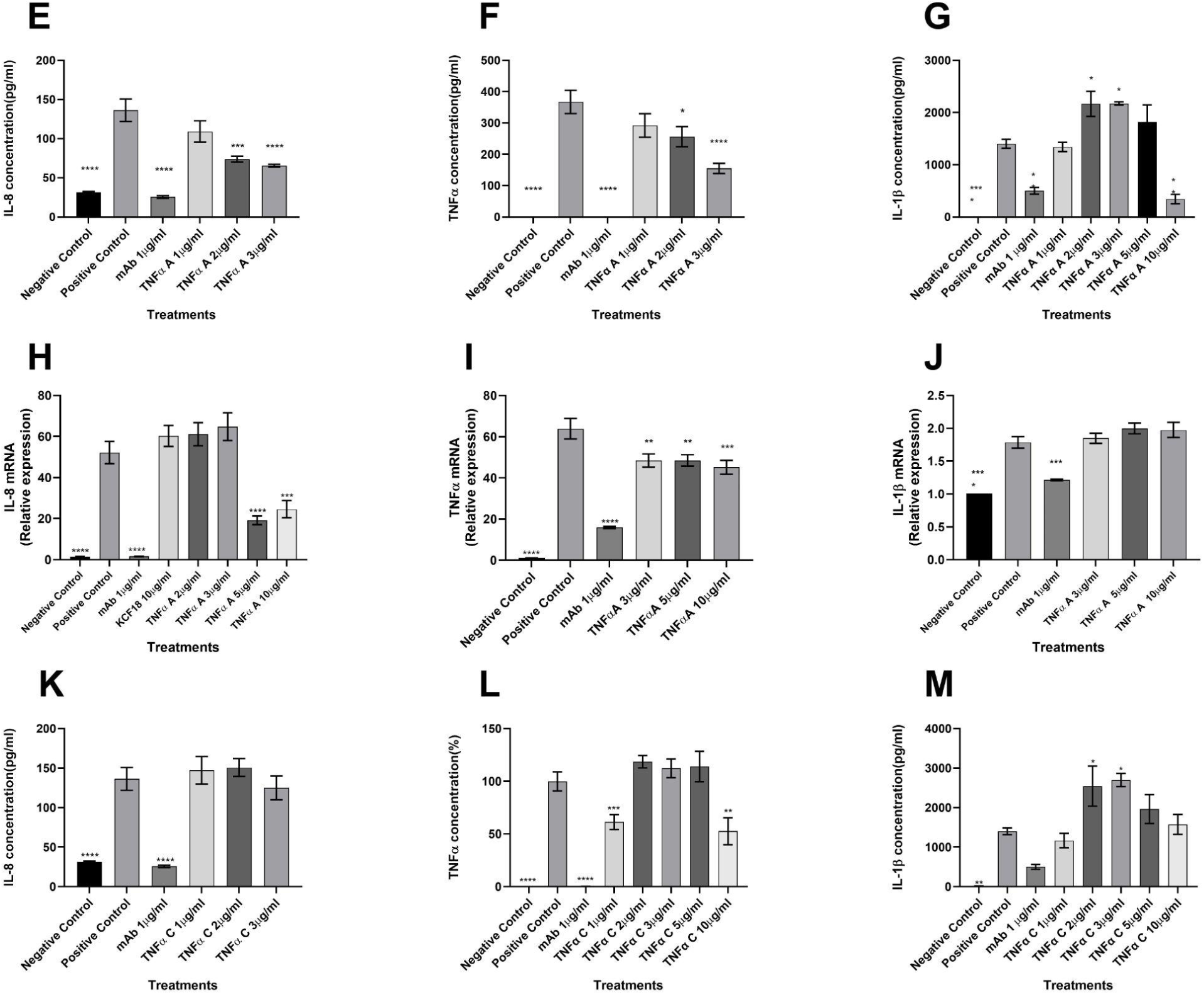
Targeting the interface between TNFα and TNFR1 using α-helical peptides. A-D computational design. 3E-J experimental measure of the: effect of increasing concentrations of the designed TNFα A peptide (AEDKVRSG) after 24 h on primed Caco-2 intestinal cells and M1-activated THP-1 cells. (K-M) Effect of addition of increasing concentrations of designed TNFα C peptide (SEDESGLG) after 24 h on primed Caco-2 intestinal cells M1-activated THP-1 cells. **(A)** Overview of TNFα - receptor interaction (homology model based on the TNFα-TNFR2 interaction 3alq): TNFα (green) forms a trimer that interacts with three separate receptor chains (one shown in cyan), each binding at a TNFα dimer interface. Only the n-terminal CRD1 and CRD2 regions of the receptor are included. Key features at the interface are highlighted: Binding pockets determined by computational solvent mapping (white mesh), Interface hotspots (sticks) determined by computational alanine scanning (yellow) and experimentally (red). **(B)** Location of designed peptides using PeptiDerive (magenta) and PatchMAN (gray and blue). **(C,D)** Details of peptide designs TNFα-A and TNFα-C designed using PatchMAN. **(E-G)** Changes in secretion, TNFα-A (**H-J)** Changes in mRNA expression levels, TNFα-A. **(K-M)** Changes in secretion, TNFα-C**. (E-M)** legend as in Figure 2.

While segments extracted using PeptiDerive (Approach A) partially covered the hotspots and pockets **(Figure 3B)**, they are unlikely to be stable (due to removal of the interactions that stabilize the derived segment within its native context of the full R1 receptor). We therefore used the PatchMAN Strategy (Approach B) to extract potential peptide backbones from interactions with a sub-domain similar to our target (see **Supplementary Figure S4** and **Table S1B for additional details**). The selected peptides form short, predicted to be stable α-helices that fill one of the potential binding pockets with either hydrophobic Leu or Ala residues and are predicted to form several salt bridges and hydrophobic interactions with TNFα **(Figure 3C,D)**. Among the suggested peptides, three peptides were selected for further study **(Table I)**. Two were further examined for activity (TNFα-A AEDKVRSG and TNFα-C SEDESGLG), while the third peptide, TNFα-B, aggregated in solution.

### Experimental validation of TNFα A peptide shows its efficiency in reducing inflammatory response

We first tested the effect of TNFα A peptide (AEDKVRSG, **Figure 3E-J**). IL-8 secretion from Caco-2 cells decreases in a dose-dependent manner upon treatment (**Figure 3E**), similarly to TNFα secretion from M1-activated THP-1 cells (**Figure 3F**). Significant inhibition was observed for both after addition of 2-3 μg/ml of the peptide. The effect on IL-1β release however did not show this pattern: at lower concentrations, the peptide led to higher (rather than lower) level of secretion, and only at high concentrations (10 μg/ml) was shown to have an inhibitory effect (**Figure 3G**). As for downregulation of mRNA expression, addition of 5 and 10 µg/ml TNFα A peptide together with rTNFα showed significantly reduced the expression of IL-8 in Caco-2 cells (**Figure 3H**), while a smaller reduction of TNFα expression in primed THP-1 cells was observed for all the tested concentrations (**Figure 3I**). Again, no effect on IL-1β expression was observed (**Figure 3J**). Thus, TNFα A peptide may significantly affect the inflammatory storm at the concentrations mentioned above.

The designed peptide TNFα C was ineffective in reducing IL-8 secretion from Caco-2 cells at all concentrations measured (**Figure 3K**; 1-3 μg/ml, tested also for 5 and 10 μg/ml, not shown). As for its effect on primed THP-1 cells, TNFα secretion was reduced at the lowest and highest concentration tested (i.e., 1 and 10 μg/ml, **Figure 3L**). TNFα C peptide was ineffective in inhibiting the release of the pro-inflammatory cytokine IL-1β, on the contrary - IL-1β release was increased upon addition of the peptide at intermediate concentrations (**Figure 3M**).

These results (IL-6 and TNFα-A peptides) suggest that both strategies, PeptiDerive as well as PatchMAN, can be used to successfully design helical immunomodulatory peptides.

### Design of β-hairpin IL-1β inhibitor using PeptiDerive

The interaction between IL-1β and its receptor subunits IL-1RI and IL-1RAcP IL1R subunits forms a “ball-and-socket” architecture, where the small globular cytokine is “engulfed” by IL-1RI and contacted by adjacent IL-1RAcP **(Figure 4A)**. We filtered PeptiDerive results for segments that overlap with identified hotspot residues and binding pockets to form stable structures (see Supplementary Materials for details). The analysis highlighted a peptide segment derived from the IL-1β cytokine (**see Supplementary Table S1A for additional details**). The peptide forms a stable β-hairpin, partially fills two potential binding pockets **(Figure 4B,C)** and recapitulates five computational hotspots. This peptide, as well as a cyclized form were selected for synthesis and characterization.

**Figure 4:**
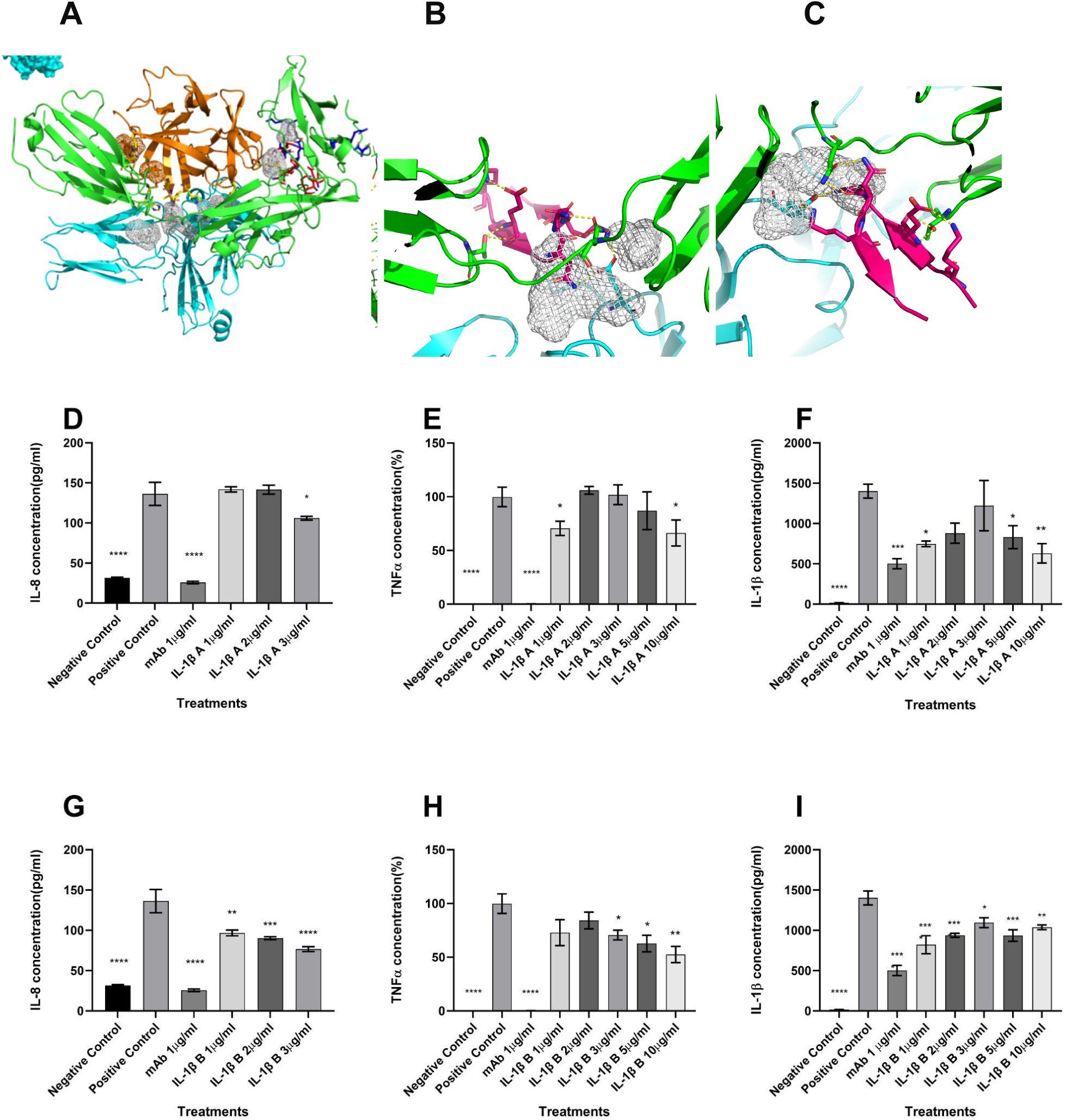
Targeting the interface between IL-1β and IL-1RI/IL-1RAcP using a β-hairpin peptide. A-C: Computational Design. D-I Experimental characterization of effect of addition of increasing concentrations of designed β-hairpin IL-1β A peptide (NKIEINNKTEE) (D-F), and its cyclized form IL-1β B (G-I), after 24h on primed Caco-2 intestinal cells and M1-activated THP-1 cells. **(A)** IL-1β (orange) binding to receptor IL-1RI and IL-1RAcP chains (green and cyan, respectively). Key features at the interface are highlighted as: Binding pockets and interface hotspots determined by computational solvent mapping and alanine scanning (as in Figure 3). **(B,C)** The inhibitory peptide generated using PeptiDerive forms a stable β− hairpin structure (magenta) and overlaps with key features of the interface. **(D-I)** legend as in Figure 2.

### Experimental validation of IL-1β β-hairpin peptides shows their efficiency in reducing inflammatory response

In Caco-2 cells, addition of the IL-1β A derived peptide (NKIEINNKTEE) moderately inhibits the secretion only at the highest concentration that was tested (3 μg/ml) **(Figure 4D)**. In THP1 cells, a significant effect was observed, but not in a dose dependent manner **(Figures 4E,F)**.

Since natural peptides are often not very stable due to their degradation, cyclization is commonly used to stabilize a peptide by reducing its flexibility, and preventing access to exoproteases. We designed and synthesized head to tail cyclic IL-1β A derived peptide. As expected, the cyclized form, IL-1β B (NKIEINNKQEE), showed improved potency and strongly reduced IL-8 secretion from primed Caco-2 cells **(Figure 4G)**, and also TNFα secretion from M1-activated THP-1 cells **(Figure 4H)**. Interestingly, the cyclic peptide also significantly reduces IL-1β secretion, but not in a dose-dependent manner **(Figure 4I)**.

In summary, peptides designed against all three cytokines tested show effective anti-inflammatory effects at low micromolar concentrations. Effects on IL-8 secretion from primed Caco-2 cells and TNFα secretion from M1-activated THP-1 cells show consistent dosage-response, where higher concentrations have a stronger effect, or an overall strong effect for all administered concentrations of inhibitor. The effect on IL-1β release from THP-1 cells seems to be more complex and requires further refinement and analysis.

## Discussion

This study presents five peptides with the potential to modulate three cytokine-receptor interactions - TNFα, IL-1β, and IL-6, all of which are fundamental in pathogenesis of IBD. In the past several decades, treatment of IBD has shifted from general, non specific anti inflammatory treatment (e.g steroidal drugs), to anti-cytokine biological therapeutics. These drugs, mainly in the form of monoclonal antibodies targeting TNFa, revolutionized the field and paved new avenues for IBD management. However, IBD treatment regimens still suffer from major caveats such as high immunogenicity, lack of response or loss of response over time, adverse effects and high logistic costs. To fill the therapeutic gap, many cytokine-aimed leads were developed in recent years, with only few proving successful.

Peptides have been shown to modulate protein interactions with efficacy competitive with antibodies, but at a much smaller molecular size. This may reduce their immunogenicity, simplify cell permeability, increase bioavailability, lower costs and reduce cytotoxicity. Many previous studies explored the potential of peptides as modulators, and several recent reports demonstrated their potential in cytokine-based systems, and specifically for multi-target modulation of cytokine networks in IBD.

In a previous study three peptides that were designed based on the structures of the interactions studied here were concatenated into one larger peptide [^27^ ^28^]. While that study suggests that this peptide (termed KCF18) reduces inflammatory response, it suffers from some crucial weaknesses: First of all, it is not clear from structural analysis how the specific selected sequence should work, since the peptides included do not cover the regions that are predicted to bind strongly. Second, the backbone of the selected peptide is not likely to maintain structural integrity when taken “out of context”. Third, the peptide used as negative control partially contains the same segments included in the binding peptide, which questions its use as negative control, and finally, measured affinity values were derived from a poor fit of non-saturated SPR curves.

In this work, we took off these starting points, and explored a scalable, straight-forward approach for peptide design for multi-target cytokine modulation. Our experimental validation suggests that the peptides that we designed “out-of-the-box”, without sophisticated sequence optimization, are effective within the low micromolar range. This further emphasizes the promise of structure-guided peptide design approaches. Given accurate identification of core interaction interfaces, natural peptide motifs such as those automatically extracted using PeptiDerive may act as peptide modulators without additional optimization. Additionally, natural structural motif templates for design may be introduced effectively, as accomplished here using a modified version of PatchMAN. With regards to IBD, our results support the hypothesis that peptides may be effective IBD modulators.

Multiple potent cytokine modulators that are both efficient and selective have recently been reported, including small molecules [^22^] and mini proteins [^21^]. Massive, high-throughput experiments were key to success for the experimental screening of small molecule libraries to identify a low-micromolar binder, and additional subsequent experiments were needed to determine the interaction key features and potential binders thereof [^22^]. The recently reported picomolar affinity inhibitory miniprotein designs targeted against IL-6 and IL-1 cytokine-receptor interactions represent an impressive and promising advance [^21^]. Of note however, this achievement relies on vast amounts of computation resources for scaffold sampling and optimization, and necessitated a large number of experimental validations.

In comparison, our work suggests a fast and straightforward strategy to first identify key interaction features and use them for a natural motif guided design, with high binder yield and dramatically less resources. Still, our work has clear limitations: Even though most of our tested peptides were successful in reduction of response to cytokine administration, the number of evaluated peptides remains small. Furthermore, some of the validation results for these peptides were inconclusive. A more thorough evaluation is required in order to appreciate the design approach and the biological system. Moreover, further binding assays and specificity tests remain to be explored. Future work should also investigate the effects of various peptide combinations, either as separate peptides or concatenated, as done for KCF18. Computationally, a more generalized scheme for evaluation of designs is needed (to systematically account for overlap with key features, predicted stability, etc). The latter is likely to benefit greatly from state-of-the-art scoring methods.

This study was conducted as the field of structural biology is being revolutionized by deep learning based methods. While the approach presented in this study is simple and low resource, we believe that the availability of new methodologies such as RFdiffusion [^43^] for de novo peptide backbones and ProtMPNN [^44^] for the sequence optimization of natural motifs can further significantly improve the inhibitory activity, stability and solubility of such peptides, as well as miniproteins.

Finally, this work presents five therapeutic lead peptides for multi-target inhibition of inflammatory cytokines in IBD, TNFα, IL6 and IL-1β. It suggests a low resource, simple to use to, effective approach for the computational design of such peptides, and insights with regards to the experimental validation of such peptides and biological systems.

## Materials and Methods

### Computational Peptide Design Strategy

Our approach aimed to identify the core structural components of interactions, and design peptides that would interfere with the interactions in these areas.

Two different strategies were used to identify peptide seeds (see **Figure 1**): A) Peptides derived from the cytokine/receptor complex (using PeptiDerive [^30^ ^29^]); and B) Peptide backbones extracted from proteins with local structural similarity to the binding pocket (using PatchMAN [^31^]).

Design steps were performed on structural models of each cytokine-receptor interaction.

1. Define interaction partners based on experimental literature - for each cytokine studied, the common partners (receptors) were identified (using UniProt [^45^], PDB [^46^] and the general literature).
2. Generate structural model of the interaction -

a. Structural data collection – solved structures of cytokine-receptor interactions were collected from the PDB [^46^]. Additionally, structures of the individual (“apo”) partners were used to infer conformational changes upon binding. For each interaction, all solved structures of either partner or the bound complex were collected. The different structures were aligned and compared for areas of flexibility or structural diversity.
b. Computational optimization of interaction models -

i. TNFα interaction model - structure of TNFα cytokine trimer (pdb 1tnf[^41^]) was superimposed onto the structure of TNFβ bound to TNFr (pdb 1tnr [^40^]) as homolog template, followed by the RosettaDock refinement protocol [^47^]. $ROSETTA_BIN/docking_protocol.mpi.linuxgccrelease -database $ROSETTA_DB -in:file:s input.pdb -native native.pdb -nstruct 350 -unboundrot unbound.pdb -use_input_sc -ex1 -ex2aro -docking_local_refine -partners [ABC_R]
3. Characterize interface and determine core components –

a. Interaction models were used to map potential binding pockets (using computational solvent mapping, FTMap [^39^], see below), and computationally scanned for hotspots (using Rosetta Alanine Scanning [^38^], see below). Experimental hotspots were collected from the literature. Additionally, published modulatory peptides for these interactions were collected to illuminate potential modulation modes.
b. All interaction characteristics were compiled and illustrated using Pymol [^48^]. Areas of extensive overlap between key features (pockets, mutations, etc) defined the core components of the interaction. See Supplementary Materials for Pymol sessions and tabular outputs. Both of the strategies employed (and described below) aim to design peptides that are able to bind the core components of the interactions, by interfering with either of the natural partners at the targeted areas. Strategy A to design inhibitory peptides consisted of the following next steps:
4. Derive short dominant peptides at the interface – short peptide segments were “trimmed” out of the binding partner (cytokine or receptor, one at a time) using the Rosetta PeptiDerive protocol [^30^ ^29^]. The derived peptides were ranked according to highest [binding energy/buried surface area] ratio. Cyclic (N-to-C termini, or disulfide bond-closed) peptides were suggested where applicable. For each “sub-interaction” (contacting a distinct region on the receptor surface) one peptide representative was selected (best rank). $ROSETTA_BIN/PeptideDeriver.linuxgccrelease -database $ROSETTA_DB -in:file:s input.pdb -peptide_deriver:pep_lengths 5 6 7 8 9 10 11 12 13 -peptide_deriver:dump_peptide_pose true -peptide_deriver:dump_cyclic_poses true -peptide_deriver:dump_report_file true -peptide_deriver:restrict_receptors_to_chains A -peptide_deriver:restrict_partners_to_chains R -peptide_deriver:do_minimize true -peptide_deriver:optimize_cyclic_threshold 0.35
5. Sequence design – derived representatives selected in the previous step were redesigned for improved solubility using Rosetta design (Fixed backbone design, fix_bb [^49^]), mainly by changing residues that previously pointed into the protein core from hydrophobic to soluble amino acids. Each such representative was inspected with respect to the key features described in the previous steps. $ROSETTA_BIN/fixbb.linuxgccrelease -database $ROSETTA_DB -s input.pdb -ex1 -ex2aro -use_input_sc -nstruct 5 -resfile design_resfile.txt -scorefile design.score.sc Example resfile: NATRO START 200 B POLAR 201 B POLAR 205 B POLAR 208 B POLAR
6. Select promising binders - representatives that recover hotspots (reported from experiments, or determined using computational alanine scanning, see step 3 above) and overlap with predicted pockets were selected for experimental validation. For cytokine TNFα, no promising peptides could be generated using strategy A. This is either due to no coverage of pockets/hotspots, or peptides that were unlikely to be stable or fold into the predicted structure (e.g. too long and lacking secondary structure). Thus, for these interactions strategy B was used. The next steps applied were as follows:
7. Map hotspots and binding pockets on the surface of the cytokines or receptors (see step 3 above).
8. Define patches of interest: for TNFα, we focused on a key interaction interface surrounding a pocket. The residues delimiting the pocket were defined as a patch (instead of automatically defining patches, as in the standard PatchMAN protocol).
9. Identify matches for the defined patch using the PatchMAN protocol.
10. Define peptide backbones (“seeds”): Use PatchMAN to extract backbone fragments of fixed lengths that are complementary to the query binding pocket, from structures in the Protein Data Bank (PDB). We used the PatchMAN design option as described in https://github.com/Furman-Lab/PatchMAN/blob/master/bin/extract_peps_for_motif.py, in which only the length of the peptide is provided (and not the peptide sequence as in docking). To allow for more flexibility in MASTER matches, we used a RMSD cutoff of 3Å (instead of 1.5Å in docking). PatchMAN_protocol.sh -c 3 <QUERY_PDB> <PEPTIDE_LENGTH>
11. Redesign the selected peptide backbones (using Rosetta FlexPepDesign, an implementation of Design into the FlexPepDock refinement steps [^50^]).
12. Select promising designs: the peptides were selected based on their coverage of the hotspots, their complementarity to the binding pocket as well as the Rosetta binding energy score.

See Supplementary Figure S4 for a scheme describing this process for the TNFα-A peptide design.

### Computational analysis of solved and modeled interactions

#### Identification of binding pockets on protein surface

We applied the FTmap computational solvent mapping approach to identify preferred binding pockets on a protein surface [^39^]. A local version of FTmap with default parameters was run separately for each partner of all the targeted interactions.

#### Identification of binding hotspot residues

We applied a local version of the Rosetta Alanine Scanning protocol (identical to https://robetta.bakerlab.org/alascansubmit.jsp) to each complex structure to identify interface hotspot residues on both partners [^38^]. Hotspot residues were defined using a threshold of DDG(complex)>1.5 REU (Rosetta Energy Units, roughly corresponding to predicted kcal/mol values).

run_ROBETTA.sh <INPUT_PDB> <CHAIN 1 ID><CHAIN 2 ID>

#### Measures used to rank designs

Designs were inspected manually, and evaluated based on several metrics:

1. For the Peptiderive scheme, we used an [energy \ peptide length] ratio (the energy contribution of the peptide segment to the interaction from which it was derived, and the derived peptide length).
2. Coverage of main binding pockets.
3. Recapitulation of both computational and experimentally reported hotspot residues.
4. Similarity to previously published modulatory peptides of the discussed interaction.

### Experimental methods

#### Peptide synthesis

Designed peptides were synthesized by Peptide Synthetics (PeptideSynthetics, Peptide Protein Research Ltd). All peptides were dissolved in ddH2O and diluted with EMEM and/or RPMI 1640 before being added to cells.

The synthesized peptides are listed in **Table 1**.

#### Cell Culture

Caco-2 Intestinal Epithelial Cells (Caco-2/HTB-37) were purchased from the American-type culture collection (ATCC, Manassas, VA). Cells were thawed and grown to confluence in Eagle’s minimum essential medium, EMEM medium containing 20% (Sigma Aldrich, USA) fetal bovine serum (FBS) (Biological Industries, Israel), 1% antibiotic antimycotic (Bio-lab Ltd, Israel), and 2mM L-glutamine, 1mM sodium pyruvate supplement. Cells were trypsinized (using 0.25%; Invitrogen, USA) and passages every 3-4 days. The medium was changed every other day.

THP-1 macrophages were provided by Prof. Gabriel Nussbaum (The Hebrew University of Jerusalem) and cultured according to specifications in his previous studies[^51^]. In short, cells were incubated in RPMI-1640 medium (ATCC and/or Biological Industries) supplemented with 10% fetal bovine serum (Biological Industries, Israel), 1% penicillin-streptomycin (Biological Industries, Israel), 10mM HEPES, 2mM L-glutamine and 1mM sodium pyruvate (Biological Industries, Israel) until they reached 70% confluence. Primary monocyte differentiation was induced following treatment with 12-O-Tetradecanoylphorbol 13-acetate (PMA) (Sigma Aldrich, USA).

The cells were plated at a concentration of ∼1×10^6^ cells/mL for 24 hours, stimulated using recombinant TNFα (rTNFα) for Caco-2 cells or LPS for THP-1 cells (see below), and treated with different concentrations of peptides. Then the cells were incubated for an additional 24 hours.

### MTT cytotoxicity experiments

MTT cytotoxicity experiments that evaluate the putative cytotoxic effect of our novel designed peptides assured that none of the peptides lead to decreased cell viability, and therefore, any decrease in the secretion or expression of cytokine mRNA levels is not due to induced cell mortality mediated by the addition of the peptide (see **Figure S1** for assays on Caco-2 cells with TNFα A, TNFα C, IL-6, IL-1β A, and IL-1β B peptides, respectively; and **Figure S2** for corresponding assays on THP1-macrophage cells). All the designed peptides tested were dissolved in ddH2O. Cytotoxic concentrations of all treatments on Caco-2 and THP-1 cells were determined using cell viability experiments using 3-(4,5-dimethylthiazol-2-yl)-2,5-diphenyltetrazolium bromide (MTT) (Sigma Aldrich USA). The cytotoxic effect was measured as cell viability percentage compared to control cells without treatment, which represent 100% cell viability.

### Calibration of concentrations for the induction of cytokine secretion

The appropriate concentration of TNFα for the induction of the maximal response for cytokines secretion in Caco-2 intestinal cells was determined by testing different TNFα concentrations (**Figure S3A-C**). Based on these experiments, we chose to use 100 ng/mL TNFα for the induction of maximal response, since lower levels did not significantly induce any significant response. Human TNFα antibody is a recombinant monoclonal human IgG1 clone used for alleviating inflammation status associated with IBD. All tested concentrations (1, 5 and 10 µg/ml) significantly reduced IL-8 secretion in response to TNFα stimulation (**Figure S3D**), and therefore, we chose to work with 1 µg/ml.

THP-1 cells were cultured to undergo differentiation in RPMI-1670 medium containing 10 ng/ml PMA for three days (as recommended for these specific cells by Prof. Gabriel Nussbaum). Upon PMA addition to the medium, the cells underwent a structural change typical of M0 macrophages and adhered to the culture plate. LPS (10 ng/ml) was then added to stimulate THP-1 to switch into a reactive, pro-inflammatory phenotype (M1). Cytokine secretion and expression was measured at several time points following the addition of LPS. For induction of maximal response, we chose to use 10 ng/mL LPS for 12h for mRNA expression and 4h for protein secretion. The expression of cytokines was measured as relative expression using Real Time qPCR and ELISA (see below).

### Calibration of positive control

Human TNFα antibody is a recombinant monoclonal human IgG1 clone used for alleviating inflammation status. The concentrations used in our system were obtained from the concentrations tested on the Caco-2 cellular system. No significant difference was observed between the three concentrations tested (1,5 and 10 µg/ml) (**Figure S3D**). According to this test, we continued to work with the concentration of the antibody in this cell system - THP-1 cells.

### Enzyme-Linked Immunosorbent Assay (ELISA)

Cells that were plated in 24 well plates were treated as described above. The medium was removed and assayed for IL-8, TNFα, and IL-1β levels using ELISA methodology according to manufacturer instructions (Invitrogen, USA).

### RNA extraction and cDNA synthesis

For THP-1, RNA was extracted using a column-based kit (NucleoSpin RNA, ORNAT, Israel). For Caco-2, RNA was extracted using an additional column-based kit (QIAGEN, RNeasy Mini kit). RNA was quantified using Nanodrop 2000 (ThermoFisher USA). 1.5 µg of RNA was used to synthesize cDNA, using qScript cDNA Synthesis Kit (Quanta bio, USA).

### Quantitative reverse transcription PCR (RT-qPCR)

Real-time qPCR was performed using Fast SYBR green master mix (ABI, USA) on Quant studio 1 machine (ABI, USA). For normalization of gene expression in all reactions, we used the GAPDH gene for TNFα, IL-8, and IL-1β gene normalization. Expression was quantified using the run standard curve method. Primers for relative gene expression are depicted in **Supplementary Table S2**.

### Statistical analysis of in-vitro experiments

All statistical analyses were performed on JMP pro 14 (SAS Institute Inc., Cary, Nc, 1989-2019) or GraphPad Prism (version 8 GraphPad Software, San Diego, California USA, www.graphpad.com). Unless otherwise stated, data are expressed as mean ± SE. Comparison between means of more than two groups was analyzed using means ANOVA and Tukey HSD.

## Author contributions

G.K.A, Z.H, B.S, T.T and O.S.F conceived the preliminary concepts. All authors participated in designing the research. Z.H, O.S.F and B.S supervised the research. T.T characterized all the cytokine-receptor interactions.

T.T computationally designed, screened and optimized all Peptiderive based designs. A.K and T.T designed screened and optimized PatchMAN based designs. G.K.A and O.G conducted all biochemical and in-vitro assays. T.T, O.S.F, G.K.A, Z.H and B.S analyzed the experimental results. T.T, O.S.F and B.S wrote the manuscript with input from the other authors. All authors revised the manuscript. O.S.F secured funding.

## Data availability statement

Models of the interactions of the designed peptides with their target cytokine/cytokine receptor are available as supplementary pdb files with corresponding names. Pymol sessions describing the different interactions and FTMap outputs, as well as outputs for computational alanine scans and peptiderive protocol are available as described in supplementary information.

## Code availability

Methods implemented in the Rosetta modeling suite (https://www.rosettacommons.org) (RosettaDock refinement, alanine scanning, Peptiderive) are available to academic and non-commercial users for free. For this study, we used Rosetta version 217 2019.14. Commercial licenses for the suite are also available through the University of Washington Technology Transfer Office. PeptiDerive is also available via an online server (https://rosie.rosettacommons.org/peptiderive). The FTmap suite is available for academic users in https://ftmap.bu.edu/serverhelp.php. For this study, we used FTMap - atlas-1.9.0-alpha12 (local). The PatchMAN protocol is available via an online server at https://furmanlab.cs.huji.ac.il/patchman/, and the code can be downloaded at https://github.com/Furman-Lab/PatchMAN/releases/tag/v1.0. Runline commands and arguments are described in detail in the Methods section.

## Acknowledgements

This work was supported, in whole or in part, by the Israel Science Foundation, founded by the Israel Academy of Science and Humanities (grant numbers 717/2017, 301/2021 to O.S.F.) and by the Clore Foundation Scholarship for PhD students (T.T).

