## Supplementary Information for "Peptide therapeutic leads for multi-target inhibition of inflammatory cytokines in Inflammatory Bowel Disease - computational design and in-vitro validation"

**Table S1 Details of peptide designs and origin of selected templates: A.** Peptides designed using peptiderive, and their respective derived natural motifs. **B.** peptides designed using PatchMAN and the selected backbone seeds.

**A**

| Designed peptide sequence | Natural motif sequence | Origin PDB, chain, positions | Uniprot ID, positions | Interaction / target | Molecular Weight (Da) |
|---|---|---|---|---|---|
| EEEQSSTRASRQ<br>IL-6 | EFLQSSLRALRQ | 1p9m, B, 174-175 | P05231, 200-211 | IL-6 – IL-6rα receptor | 1407 |
| NKIEINNKTEE<br>IL-1β A | NKIEINNKLEF | 4dep, A, 107-117 | P01584, 218-228 | IL-1β – IL-1R receptor | 1331 |
| NKIEINNKQEE<br>(cyclic peptide)<br>IL-1β B | NKIEINNKLEF | 4dep, A, 107-117 | P01584, 218-228 | IL-1β – IL-1R receptor | 1358 |

**B**

| Designed peptide sequence | Bacbkone seed sequence | Origin PDB, chain, positions | Uniprot ID, positions | Interaction / target | Molecular Weight (Da) |
|---|---|---|---|---|---|
| AEDKVRSG<br>TNFα A | YVDLRDKM | 1hto,A, 194-201 | P9WN39, 195-202 | TNFα-TNFRI cytokine | 861 |
| SEDESGLG<br>TNFα C | PFIRSHVR | 3hrd,C, 84-91 | Q0QLF4, 84-91 | TNFα-TNFRI cytokine | 793 |

**Table S2 Primers for measure of relative human gene expression**

| Name | Forward 5' □ 3' | Reverse 5' □ 3' |
|---|---|---|
| GAPDH | TCACCAGGGCTGCTTTTAAC | GACAAGCTTCCCGTTCTCAG |
| TNFα | ATCTTCTCGAACCCCGAGTG | ATGAGGTACAGGCCCTCTGAT |
| IL-8 | AGTCCTTGTTCCACTGTGCC | GTGCTTCCACATGTCCTCAC |
| IL-6 | CCTTCCAAAGATGGCTGAAA | CAGGGGTGGTTATTGCATCT |
| IL-1β | CTGTACCTGTCCTGCGTGTT | AGACGGGCATGTTTTCTGCT |

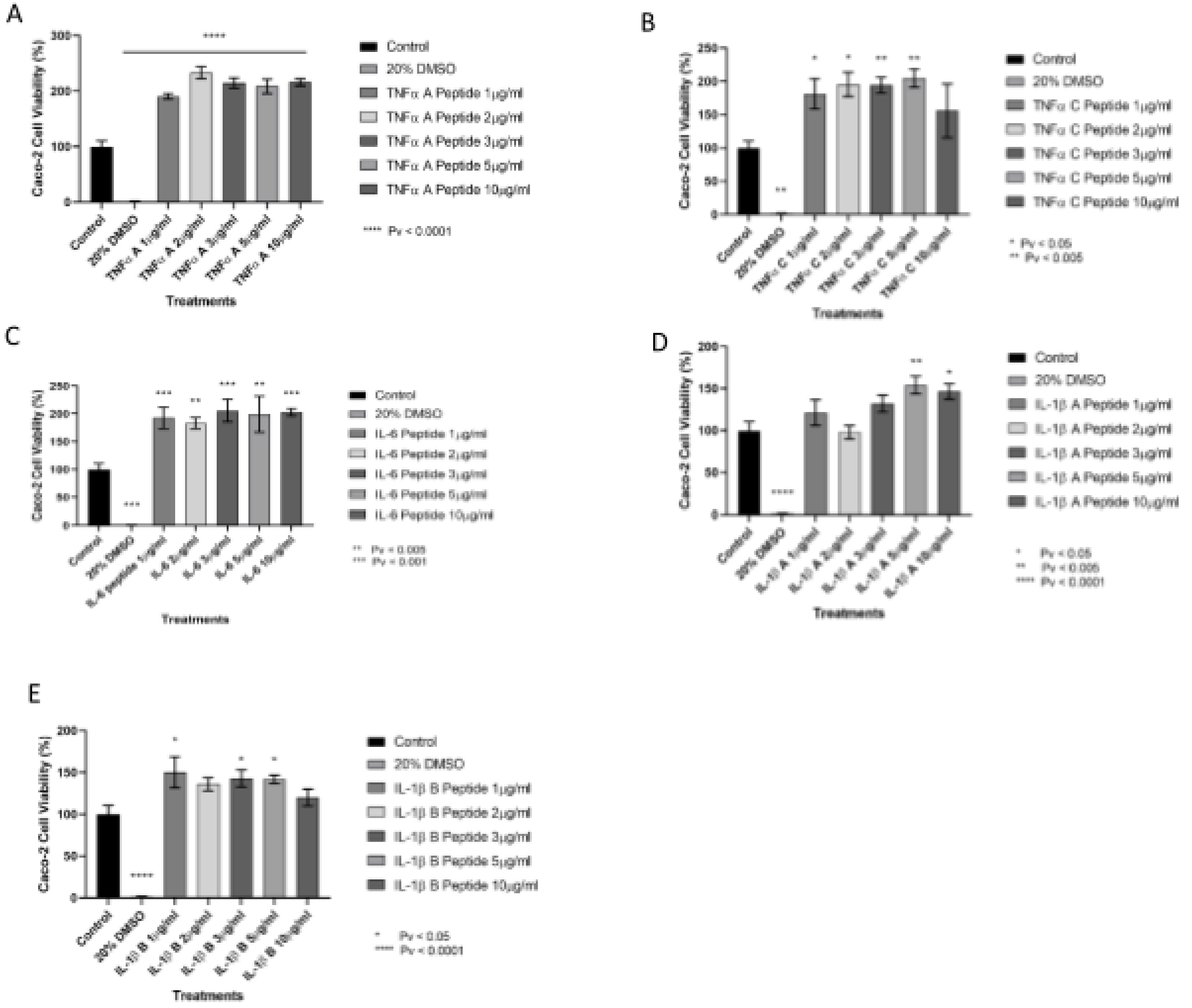


**Figure S1 MTT experiment on Caco-2 intestinal epithelial cells with different peptide concentrations**. **(A)** TNFα A **(B)** TNFα C **(C)** IL-6 **(D)** IL-1β A **(E)** IL-1β Control represents Caco-2 cells without treatment, and 20% DMSO is the negative control. Caco-2 cells were treated with 1µg/ml to 10µg/ml of the different peptides for 24 hours, n=4.

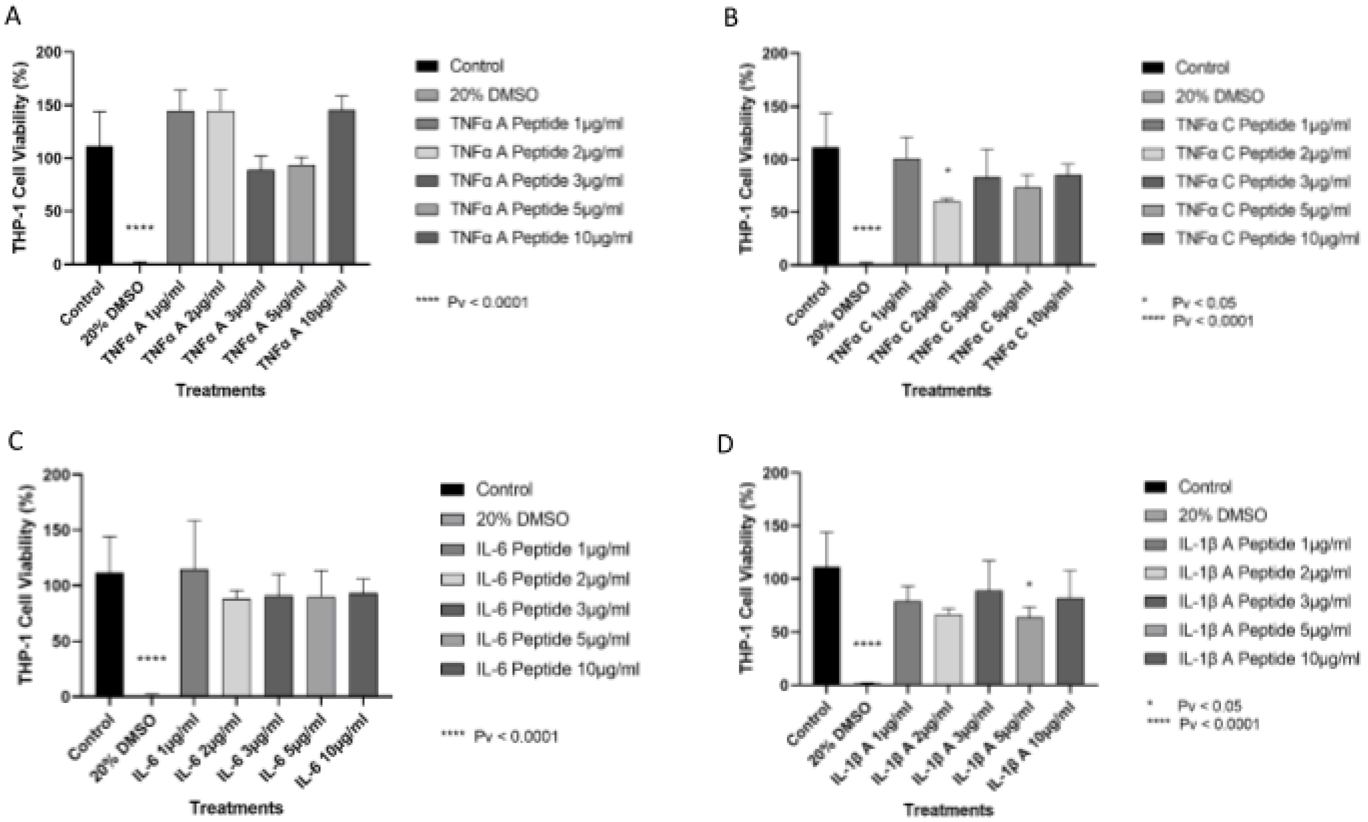


**Figure S2 MTT experiment on THP-1 macrophage cells with different peptide concentrations. (A)** TNFα A **(B)** TNFα C **(C)** IL-6 **(D)** IL-1β A, Control represents THP-1 cells without treatment, and 20% DMSO is the negative control. THP-1 cells were treated with 1,2,3,5, and 10 µg/ml of the peptides for 24 hours, n=4.

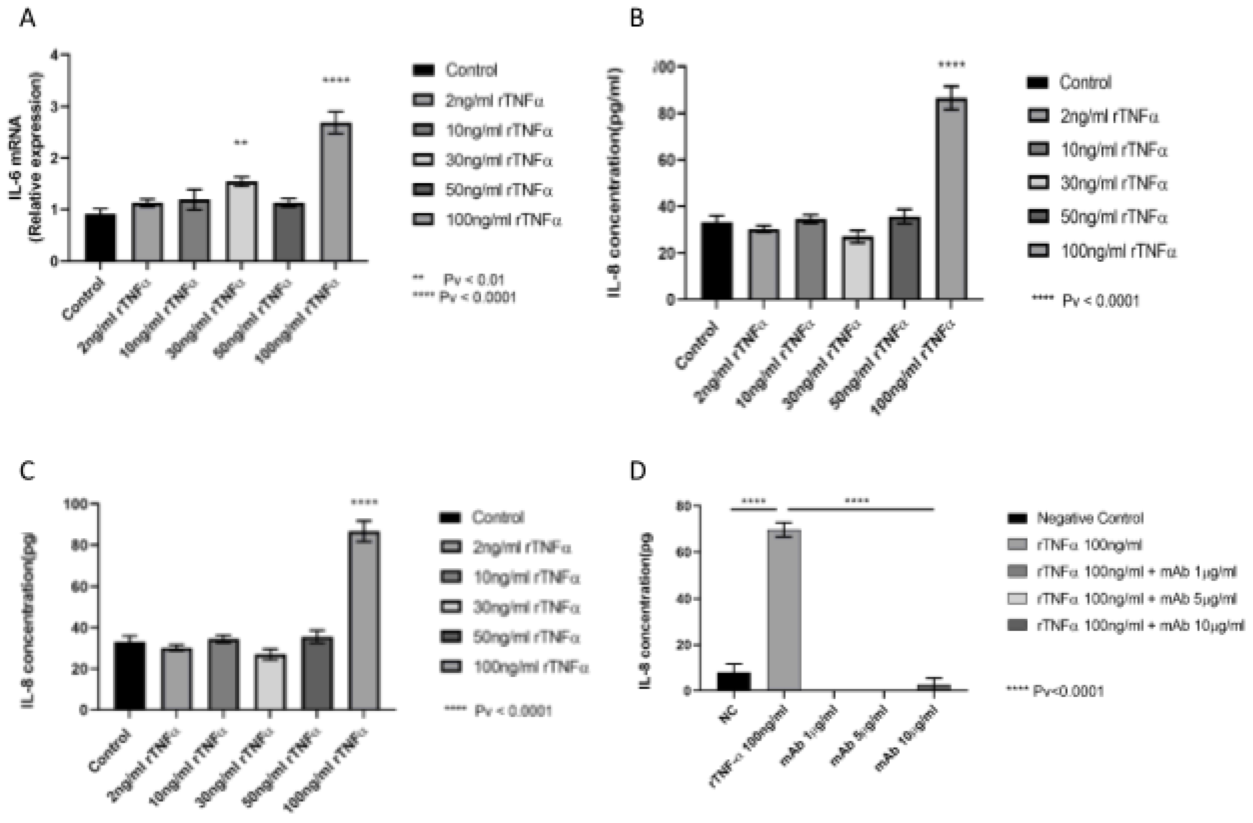


**Figure S3 Calibration of TNFα concentration on Caco-2 epithelial cells plated on 12 well plates.** (**A,B)** IL-6 mRNA and IL-8 protein levels were measured after 4 and 24 hours, respectively. **(C)** IL-8 protein level was measured 24 hours after treatment with rTNFα. (**D)** Calibration of human TNFα antibody concentration (mAb) for optimal reduction of IL-8 secretions. n=3, P <0.0001.

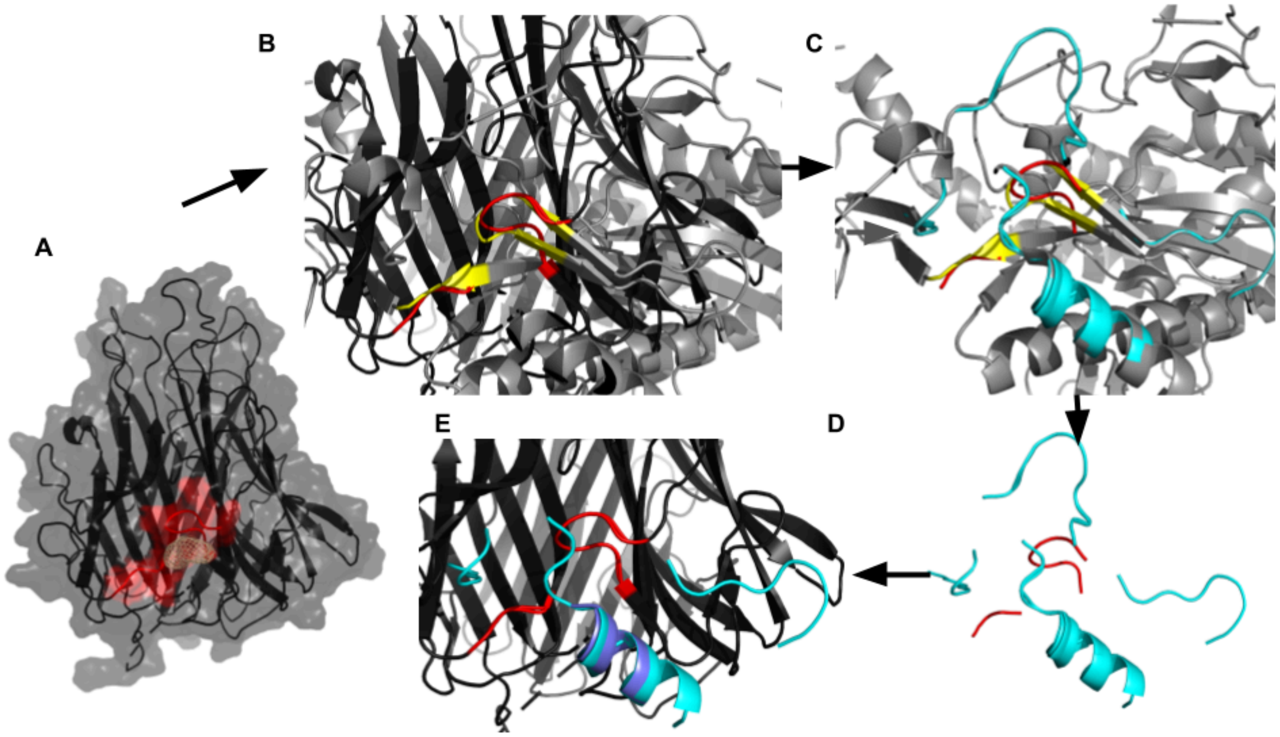


**Figure S4 Illustration of the PatchMAN design process for peptide TNFα-A. (A) TNFα interface of interest:** TNFα cytokine trimer (black transparent surface and cartoon). FTMap pocket shown as wheat colored mesh. The query for PatchMAN are residues defining the pocket colored red (cartoon and surface). **(B) Detection of structural matches:** TNFα trimer superimposed onto 1hto (gray cartoon), based on the match between query (TNFα, red) and hit (1hto, yellow), according to MASTER search, as implemented in the PatchMAN protocol **(C) Definition of potential surrounding peptide seeds:** selected backbone fragments (cyan) in close interaction with the hit residues (yellow). **(D) Extraction of backbone seeds. (E) Superposition of backbone seeds onto query protein:** The backbone selected as seed for the TNFα-A peptide design is colored slate-blue, other backbones are colored cyan.

**Additional Supplementary Material:**

Zip file containing:

1. Designed peptide models (pdb files)
    a. peptide_il6_EEEQSSTRASRQ.pdb
    b. peptide_tnfa_A_AEDKVRSG.pdb
    c. peptide_tnfa_C_SEDESGLG.pdb
    d. peptide_il1b_A_NKIEINNKTEE.pdb
    e. peptide_il1b_B_NKIEINNKQEE.pdb

2. Alanine scanning results (csv files) -
    a. il6_interaction_1p9m_alascan_results.csv
    b. tnfa_interaction_1tnf_1tnr_alascan_results.csv
    c. il1b_interaction_4dep_alascan_results.csv
3. PeptiDerive results (txt files) -
    a. formatted_il6_interaction_1p9m_cytokine_peptiderive_results.txt
    b. formatted_il6_interaction_1p9m_receptor_peptiderive_results.txt
    c. formatted_tnfa_interaction_1tnf_1tnr_cytokine_peptiderive_results.txt
    d. formatted_tnfa_interaction_1tnf_1tnr_receptor_peptiderive_results.tx
    e. formatted_il1b_interaction_4dep_cytokine_peptiderive_results.txt
    f. formatted_il1b_interaction_4dep_receptor_peptiderive_results.txt
4. Pymol sessions (including view of interactions with selected peptides, peptiderive results, Ftmap results, alanine scanning results and experimental annotation) -
    a. il6_1p9m_full_analysis_overview_session_supplementary.pse
    b. tnfa_full_analysis_overview_session_supplementary.pse
    c. il1b_4dep_full_analysis_overview_session_supplementary.pse

- File names denote the interaction being modeled (e.g. il1b and its receptor), the pdb file of the structure used for modeling (e.g. 4dep), and the type of results (e.g. alascan \ peptiderive \ overview_sessions)
- For peptiderive results, file name indicates whether peptides were derived from the receptor or the cytokine.
- Pymol sessions include:
    - Cytokine-receptor interaction
    - FTMap solvent mapping pockets ("probes" mapped to both the receptor and ligand and annotated accordingly, viewed as colorful meshes)
    - Hotspot residues - either computationally mapped (alanine scanned, yellow) or experimental (colorful, annotated according to the relevant source)
    - Derived peptides - annotated according to which partner was derived (either the cytokine or the receptor), length of peptide and whether its linear or cyclic

- Selected peptide designs are annotated accordingly, with several relevant stored views matching figures in the paper and others
